# Rapid discovery of novel prophages using biological feature engineering and machine learning

**DOI:** 10.1101/2020.08.09.243022

**Authors:** Kimmo Sirén, Andrew Millard, Bent Petersen, M Thomas P Gilbert, Martha RJ Clokie, Thomas Sicheritz-Pontén

## Abstract

Prophages are phages that are integrated into bacterial genomes and which are key to understanding many aspects of bacterial biology. Their extreme diversity means they are challenging to detect using sequence similarity, yet this remains the paradigm and thus many phages remain unidentified. We present a novel, fast and generalizing machine learning method based on feature space to facilitate novel prophage discovery. To validate the approach, we reanalyzed publicly available marine viromes and single-cell genomes using our feature-based approaches and found consistently more phages than were detected using current state-of-the-art tools while being notably faster. This demonstrates that our approach significantly enhances bacteriophage discovery and thus provides a new starting point for exploring new biologies.

## INTRODUCTION

Prophages are bacteriophages integrated into bacterial genomes where they play an important role in the ecology, physiology and evolution of their bacterial hosts(1, 2). Despite the many and varied examples of ways that prophages impact bacterial biology, our knowledge is based on a somewhat limited number of prophages in well-studied prokaryotes(3–5).

To comprehensively understand the importance of prophages and further interrogate their multitude of roles, it is first necessary to predict their presence within prokaryotic host genomes. Current bioinformatics prediction tools largely rely on sequence similarities and therefore struggle to identify novel motifs with no close analogs in the public searchable databases. The current most widely used and very useful prophage prediction tool is PHASTER(6), which carries out sensitive comparisons to existing phage genes by combining sequence similarity searches with gene presence and synteny. When it comes to extracting viral sequences from (meta)genomic data, again there is a reliance on well-understood phages and the available tools are dependent on linking the reference viral genome database with sequence similarities approaches as is the case with VirSorter(7)and VIBRANT(8).

Although some feature-based approaches have been applied to predict prophages and phages from (meta)genomic datasets (9, 10), they have either been trained on a limited set of biological features (e.g. only nucleotide frequencies), or inadequate amount of training data. Nevertheless, when applied alongside careful curation of training data, such approaches yield valuable new information, as demonstrated through the recent characterization of inoviruses(11), and significantly reduce prediction times, improving the speed and scalability of such models to big data.

To overcome the current limitations and increase our knowledge of phage space we present PhageBoost, a bioinformatics machine learning tool for fast, generalizable and explainable detection and discovery of prophage regions. PhageBoost disentangles the viral signal from the host background by shifting from sequence space into biological feature space. As proteins with similar functions can share attributes, or features, despite being far apart in sequence space(12), this makes predictions less prone to sequence similarity limitations.

Our approach calculates and engineers biological features from both nucleotide and amino acid sequences for every gene, and then uses machine learning to predict which gene belongs to bacteria or phages. The resulting prediction probabilities are parsed to longer regions, which are considered viral if their probability distributions differ from the background. PhageBoost utilizes biological features such as GC-content, amino acid composition, gene length, gene direction, intergenic distances and codon adaptation index (full list of features in Table S1), and extreme gradient boosting (XGBoost)(13) to learn the differences between the host and phage genes in relation to the complete genome signal.

In order for a machine learning model to be able to predict if a region in a bacterial genome is a prophage, the model needs to be trained on (i) a trusted, preferable experimentally verified, positive dataset of known prophage regions, (ii) a similar trusted negative dataset of strictly bacterial regions, and (iii) further validated with a similar dataset that was not used in any part of the training. Here we used all completely sequenced bacterial and archaeal genomes from NCBI (Table S2) and defined prophage regions and prophage-free regions by scanning for prokaryotic virus orthologous groups (pVOGs)(14) and clusters of orthologous groups (COGs)(15) searches respectively and keeping continues stretches of 10 genes or more. For each gene in the resulting 997,443 phage regions and 1,806,720 phage-free regions, we transformed the nucleotide sequences into 1,587 different biological features relative to the genome differences and selected 208 features that were present and highly variant throughout the training data genomes (Table S1).

As not all bacterial genomes were equally present in the training data, this could potentially skew the machine learning algorithm towards prophage regions from the most abundant bacteria. To give equal importance to less occurring bacteria, we computed sample weights for all genes by assigning them to 23,329 groups based on sequence identity and genome taxonomy. A gradient boosting decision tree model(13) was trained on 3,672,101 genes with manually fine-tuned hyperparameters to limit false positives while optimizing the logarithmic loss. We stopped the boosting rounds after no error rate decrease was observed for a test set of randomly chosen 23,329 genes, one from each group.

## MATERIAL AND METHODS

### PhageBoost workflow

For a (meta-)genome prediction, PhageBoost starts working after the gene calling by expecting nucleotides, amino acids, and coordinates as inputs for each gene. We have currently PhageBoost will start from fasta-file and implements gene prediction using cythonised Prodigal(16) from Pyrodigal v.0.2.1 (https://github.com/althonos/pyrodigal). However, any gene caller results can be used as input. PhageBoost will calculate the biological features for each gene and transform them relative to the background of the set of contigs before predicting. This is done by standardizing the features by subtracting the mean and scaling to unit variance before the probabilistic classification of each gene as phage or bacteria. Afterward, the genes above a probability threshold (default 0.9) are parsed to regions using multiple adjustable parameters that allow customizable pattern matching: length of a minimum number of genes (default 10), the neighboring genes required to have the same threshold (default 0) and the allowed gap between genes (default 5). We further smooth the predictions using Parzen rolling windows(17) of 20 periods and look at the smoothed probability distribution across the genome. We disregard regions having either a summed smoothed probability less than 0.5, or region less than one unit of standard deviation away from the negative predictions smoothed average, and finally, we reject those regions whose the probability distributions differ from the probability distribution generated from the negative predictions by using Kruskal–Wallis rank test (default, alpha: 0.001) as implemented in Scipy(18). The algorithm returns the predicted probabilities, smoothed probabilities, and predicted regions for each gene. Ultimately, each phage region can then be filtered out from the input fasta-file using a start and stop coordinates.

### Feature generation

We calculated 1,587 different features (Table S1) using the BioPython SeqUtils ProteinAnalysis module(19) and in-house scripts for each gene. To avoid model bias and make the model more generalizable, we applied simple feature engineering and selection. We selected the 208 features that were always present and high variance throughout the training data genomes. We further transformed the feature values relative to the genome differences.

### Training dataset generation

For a machine learning model to be able to predict if a region in a bacterial genome is a prophage, the model needs to be trained on (i) a trusted, preferable experimentally verified, positive dataset of known prophage regions, (ii) a similar trusted negative dataset of strictly prokaryotic regions, and (iii) further validated with a similar dataset that was not used in any part of the training. In essence, to make sure the dataset is valid, it should consist of independent and identically distributed random variables. As no golden standard dataset is currently available, we constructed a dataset which should ideally be as close as possible to reality. For the training dataset generation, we used all completely sequenced bacterial and archaeal genomes from NCBI’s RefSeq(20) up to February 2019 and chose those chromosomes which had 300 genes or more, resulting in 13,994 genomes (Table S2). In order to create a classification value for the training data, phage regions and phage-free regions were defined using labels generated from clusters of orthologous groups (COGs)(15), and prokaryotic virus orthologous groups (pVOGs)(14). The genomic regions that were designated as phage regions consisted of regions where only pVOGs exist throughout a stretch of ten genes. In contrast, the phage-free regions were defined as regions where only COGs exist throughout the ten genes stretch without the presence of any pVOGs. From the 13,994 genomes used for the model training and test data, we extracted 31,973 phage regions and 101,747 phage-free regions totaling 4,007,643 genes.

### Training sample weights

We used the individual genes found in the regions to train the classifier model. As not all bacterial genomes were equally present in the training data, this can skew the machine learning algorithm to towards prophage regions from the most abundant bacteria. Additionally, this can cause the machine learning model to learn patterns driven by particular taxonomical lineage or gene homology. To give equal importance to less abundant bacteria, and in order to limit model bias caused by redundant genes, we calculated clusters to generate sample weights during training. MMseqs(21) and MCL(22) (inflation parameter set to 2) were used to assign the genes to the gene clusters in a similar fashion as previously described(23). We applied majority voting to eliminate model bias caused by having the same features assign to both phage and prokaryote. Thus to limit the bias caused by gene clusters without the majority group, we removed the gene cluster when the dominant group proportion was less than 0.7 and pooled the gene clusters with less than ten members for more even sample weight generation. We managed the potential bias caused by taxonomy grouping genes belonging to the same family if more than 5,000 genes. From 116,512 controlled gene clusters, we generated 23,329 labels for sample weights.

### Training of the model

After filtering steps during the sample weight generation of the 4,007,643 genes, we trained the model using 3,672,101 genes as the training data. We used a test dataset of 23,329 genes for early stopping during training. These were generated by random sampling of single genes from each gene cluster. After taking a sample from each group as test data for early stopping of model training, the sample weights used in the were computed using the balanced mode from Scikit-learn v0.22.1(24). The datasets are available as supplementary data. The final gradient boosting decision tree model using XGBoost v.1.0.2(13) was trained on 3,672,101 genes until no further model improvement was observed for ten boosting rounds of the test data using classification error as the evaluation metric. The model hyperparameters were manually fine-tuned to avoid generating false positives while driving the log-likelihood score lower after getting the initial idea of parameters through a ten-fold cross-validated search with Bayesian optimization framework Optuna v. 0.9(25).

For benchmarking PhageBoost for the 54 genomes, we removed these genomes from the training data and recalculated the sample weights but used the same hyperparameters. This resulted to 3,655,262 genes in the training dataset and 23,278 clusters and 23,278 test genes for stopping the boosting (data available at supplementary data).

### Model explanations

Explaining the model predictions was done by utilizing the Shapley additive explanations for ensembles of trees from shap v.0.35.0(26) and the builtin method for current XGBoost versions for easy access to the same feature contributions and interactions for the predictions. To understand how the model learned during the training, we computed and visualized the feature contribution for the training data, as well as the link between the Shapley value and the feature value in Figure 1e and Figure S1 and individual figures and raw Shapley value data in supplementary data. We used the predicted feature contributions and extracted sorted order of the feature importance on models local output by using the standard approach taking the averages on the absolute values of the Shapley values. For Figure 1E, the feature value and Shapley value interaction were simplified using the Pearson correlation coefficient, while the figures in supplementary data have raw data visualized in a scatter plot. See Figure S1 for the barplot of the impact of each feature during training. For figures 1B and 1C and 1D, we smoothed the predictions by using Parzen(17) window rolling averages of twenty periods. We summed the raw contributions for each gene followed (1C), and visualized the smoothed ten features with the most contribution in the training dataset (with the highest absolute average) (1D), and summed total features followed by smoothing (1B). The workflow on how to create the images in figure 1 is found in the jupyter notebooks in GitHub: https://github.com/ku-cbd/PhageBoost/tree/master/notebooks.

**Figure 1.**
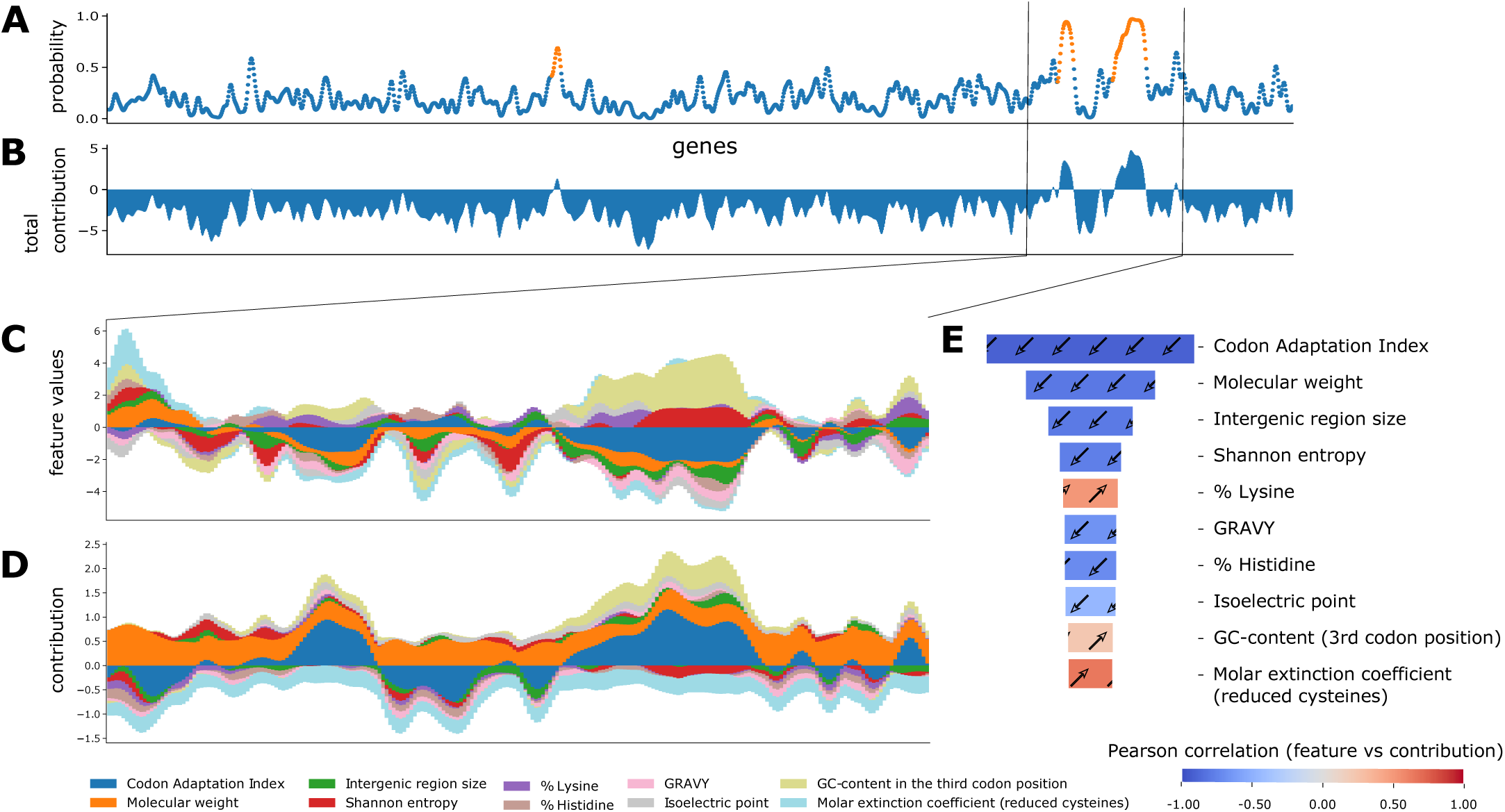
PhageBoost prophage predictions (in orange) along a bacterial chromosome (*Haemophilus influenzae*, NC_000907.1) (blue) (panel A) with total feature influence (panel B). The biological feature content varies among the prophage regions and from the bacterial genome (Panel C). Individual feature contributions can be used to explain the model predictions (Panel D)(26). This allows the extraction and validation of biological signals. Panel E shows the ten most important features learned during the training phase that the model uses to discriminate between prophage and bacterial regions. Panels C and D show the influence of the same ten features along the predicted region. Colorbar: Pearson correlation coefficient between the feature values and Shapley values.

### Benchmarking and comparisons 54 genomes

We took the approach of using the previously reported 267 prophages from 54 genomes(27) that have been previously used to benchmark prophage prediction tools(6, 7, 28). We extracted the list of the validated prophages from the PHASTER website (table 4, https://phaster.ca/statistics). For each genome in the 54 validation set for all the predictors, we used the regions begin and end coordinates (base pairs) to count the sensitivity and positive predicted values (PPV) as previously suggested for phage dataset validation(6). The sensitivity and positive prediction value (PPV) were defined as follows:

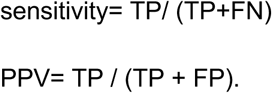

Where true positive (TP) equals the nucleotide found in both validation set and with the prediction tool, false positive (FP) equals nucleotides found only in predictions; false negative (FN) nucleotides found only from validation. However, we would like to point out, as already noted in 2016 by the developers of PHASTER, “.. given that the ‘gold standard’ annotations used for evaluation are […] old, many prophages identified as ‘false positives’ relative to this standard are likely to be true prophages.” (6). More than the reported prophages as in the genomes would negatively influence the PPV score(6, 28). To link the prediction performance for each phage region and take into account the different sizes of the viral regions, we also calculated the proportion of region retained. We generated in-house python scripts to compute all three values.

We benchmarked PhageBoost against VirSorter(7), VIBRANT(8), and PHASTER(6). We used the default settings for PhageBoost v.0.1.2, VIBRANT v.1.2.1, and VirSorter v.1.0.5 with the database db2, while we manually submitted 54 genomes to the PHASTER web server using their URLAPI (https://phaster.ca/instructions) and chose not to use precomputed results. Results have been deposited to supplementary data.

### Virome mapping

We mapped the short reads from 223 published marine viral metagenomic samples(29) to the 5539 single amplified genomes(30) Table S4. These virome samples come from 65 Tara stations. We used the Anvi’o v.6.1(31) platform for the work. We merged all the SAGs to a multi-fasta file, after which we mapped each short-read sample to the concatenated SAGs file using Bowtie2 v.2.3.5 (32) with the -a flag to return all the possible matches and otherwise default parameters to avoid signal dilution. The read recruitment varied from 0 to 5.22 percent (Table S5). Afterward, by using the gene calling done by the original authors of the dataset(30), we extracted the coverage and detection information for each gene (supplementary data). Using the detection information, we extracted the regions found by the viral mapping with a custom in-house script. We defined a region found with a minimum number of 5 genes with detection of 1.0. We parsed the regions for each virome sample separately (supplementary data). We additionally extracted the prophage-like regions from the SAGs by selecting the regions that were at least 10kb long and located 10kb away from the edges.

### Predicting viral signal from the SAGs

For the dataset of 5,537 single-amplified genomes (SAGs)(30), we used the default settings for PhageBoost v.0.1.1 predictions. These are minimum region length 10, 5 allowed gaps, and probability threshold 0.9. We used the default settings for VirSorter v.1.0.5(7) predictions with the database db2 using all the phage hidden Markov models (HMMs) and curated HMMs. For the PHASTER(6) predictions, we submitted genomes to the PHASTER web server using their URLAPI (https://phaster.ca/instructions), and thus could not be benchmarked in the same way as other tools in terms of time. And chose and chose not to use pre-computed results together with a multi-fasta option. We used the default settings for VIBRANT v.1.2.1(8). To link the predictions to the regions found by viral read recruitment, and take into account the different sizes of the viral regions, we calculated the proportion of region retained.

### Ecological significance

We further wanted to investigate the ecological significance of the filtered dataset regions found by linking this to the prediction tool. We took the metadata associated with the isolation and sampling locations for both the viromes and the SAGs and generated the data for each potential prophage region (Table S4). Using kepler-gl v.2.2.0 (https://kepler.gl/), we visualized the sites by latitude and longitude metadata of the sampling spots of both viromes and SAGs for each prophage prediction tool (Figure 3). An interactive map is provided in supplementary data. For VIBRANT, the six SAGs came all from a single sampling location; these were mapped to 121 virome sampling sites. PHASTER found seven unique locations for SAGs and 67 virome locations, whereas VirSorter found five locations for SAGs and 124 locations for viromes, and PhageBoost had most considerably more ecological signals with 11 locations for SAGs and 125 locations for viromes (supplementary data). These results suggest that some of the prophages might be more globally present than previously understood using the current tools.

**Figure 2.**
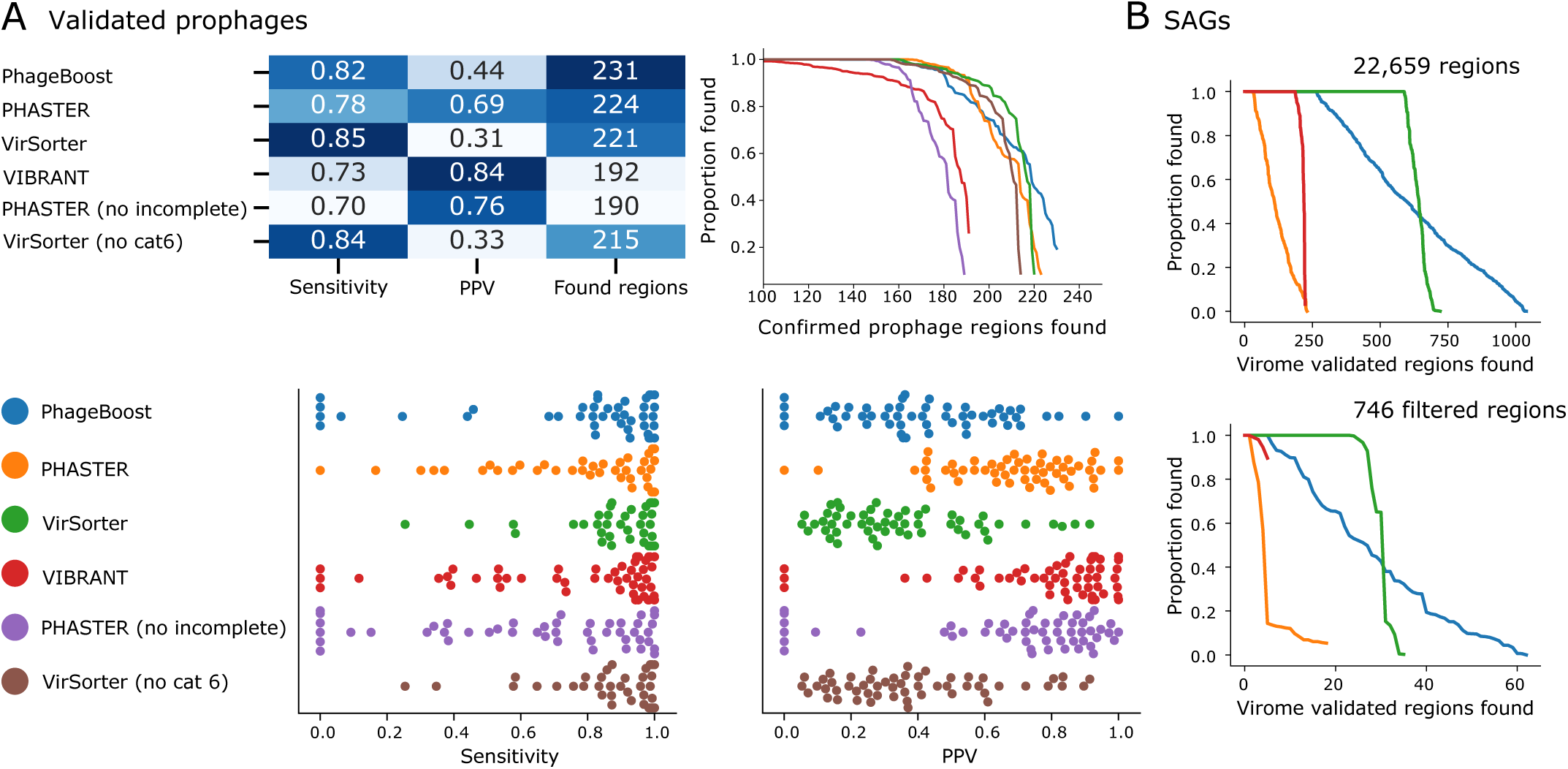
Software benchmarking. Validation against a previously reported dataset of prophages from 54 prokaryotes(27) with a total found regions, positive predictive value (PPV), and sensitivity measured for single genomes as well as phage proportion (in bp) found per prophage. (Panel A). Virome(29) mapping validated phage signals marine single-amplified genomes (SAGs)(30) and a filtered subset with a prophage-like pattern. (Panel B).

**Figure 3.**
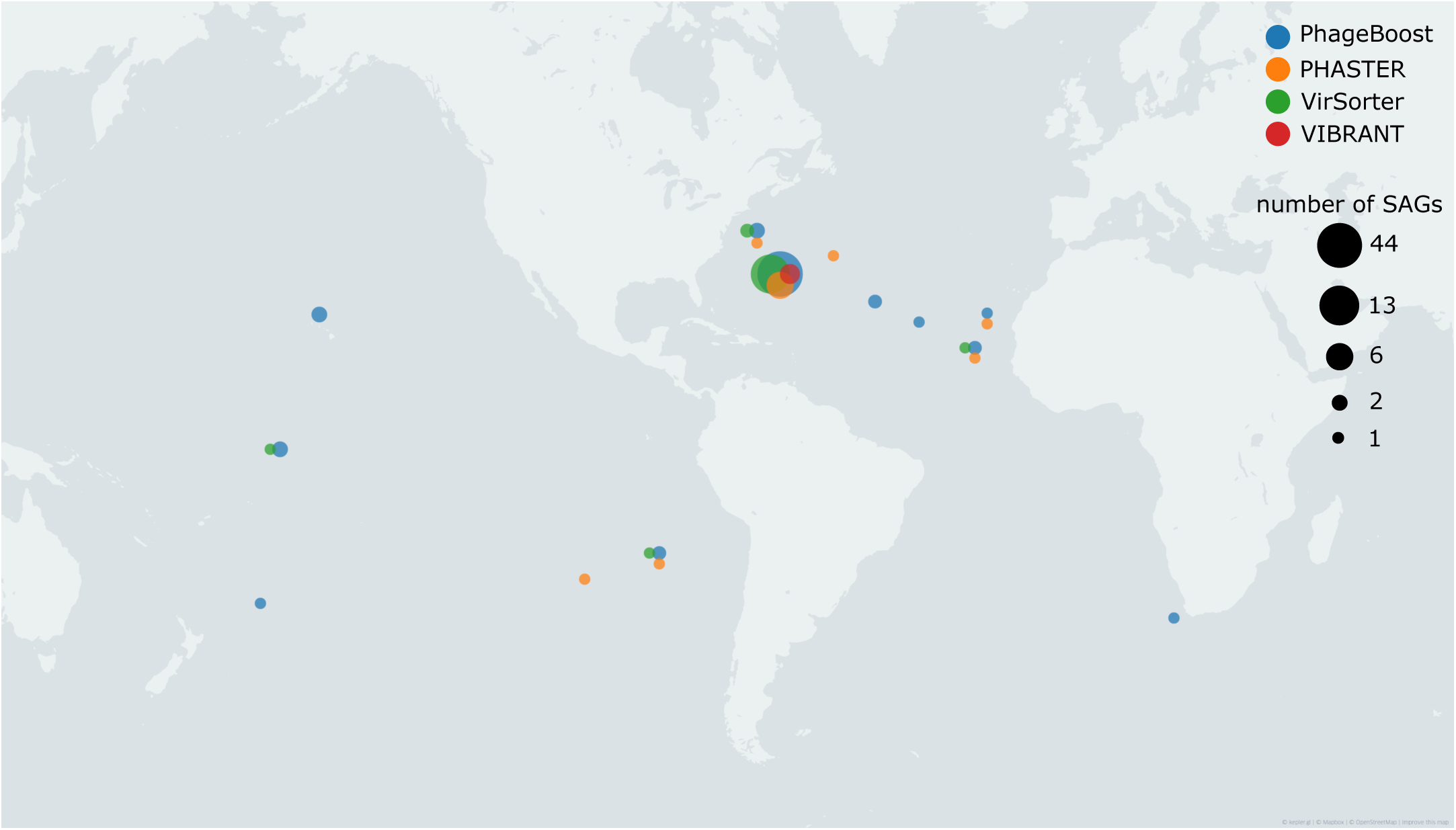
The sampling locations of the single-amplified genomes (SAGs) with potential prophage regions found by different prophage predictors and confirmed by virome reads. The coordinates are jittered if multiple prophage predictors overlap. The size of the point is dependent on the number of SAGs found from those coordinates.

We used both GNU parallel v20200322(33) and joblib v.0.14.1 to speed up the computation throughout the model preparation, parsing, and processing data for this paper. All the data visualization and figures were generated using either kepler-gl, Matplotlib(34) or Seaborn(35) and were manually fine-tuned for publication using Inkscape (http://www.inkscape.org/).

## RESULTS

### Biological explanation of discriminative features

To evaluate the model behavior, explanations of the individual predictions of the machine learning model’s outputs were created based on Shapley additive explanations(36). This approach allows an assessment of the general feature importance generated during model training and the subsequent exploration of which interactions inform predictions. Thus, we can then relate biological significance to the key feature contributions identified in our models.

To distinguish between prophage or bacterial origins, PhageBoost generates prediction probabilities for each gene across the bacterial genome or metagenomic contig (Figure 1). Of the many hundreds of biological features used, it is often smaller subsets drive the prophage prediction (Figure 1, panels B-D). During the training phase, by explaining(26) the predictor local output, we can identify subsets of discriminating features which are shown in the order of relative importance in Figure 1, panel E. A key strength of our approach is that the feature contribution can be related to the actual feature values in order to extract a biological signal that defines the prediction (Figure 1, panel E).

Although regression tree boosting models are more complex than linear univariate models, biological insights can be gained from the ranking of the feature importances. The strongest feature, which PhageBoost uses to discriminate between prophage and bacterial regions, is the Codon Adaptation Index (CAI), which measures the synonymous codon usage bias for genes with respect to a set of reference genes. Originally, the CAI is based only on highly expressed genes, but here we used the genome data as we didn’t have expression data(37). Temperate phages generally adapt their codon usage to be concordant with their hosts with time, as they utilize the host translational machinery. There is evidence for this in a small set of coliphages that have been examined, with greater adaptation in temperate phages compared to virulent phages(38). However, adaptation is further complicated by the phage carriage of tRNAs that may reduce the need to evolve codon usage. Why a low CAI is such a strong feature is not immediately obvious, it may be that the identified phages are very recent acquisitions and did not have time to evolve. Other relevant features are the length of genes and the lengths of the regions in between the genes in the observed region - which is not surprising as phage genes tend to be shorter than bacterial genes and phage genomes are more compact, thereby having shorter intergenic distances. Other important features consist of the GC content at the 3rd position in the codon, the molecular weight, and the percentages of threonine, histidine and cysteine, where more cysteine and histidine residues are signatures of prophages, and higher contents of threonine are attributes of bacterial regions. The Shannon entropy has already been shown to be discriminative between bacterial genomes and phage genomes(39) and has previously been used for detecting prophages in bacterial genomes(9). The grand average index of hydropathy (GRAVY) which is essentially a measure of hydrophobicity of proteins, and it had been shown that phage proteins used in phage display may be more hydrophobic(40). For a full set of feature importances see Figure S1 and Table S3.

### Benchmarking and validation of novel predictions

We chose to benchmark PhageBoost in two ways (Figure 2) - by comparing its performance to existing state-of-the-art methods and by discovering previously unknown viral signals by mapping marine viral fractions on to single amplified marine microbial genomes.

#### Experimentally verified prophages from 54 prokaryotic genomes

We used previously reported 267 prophages from 54 prokaryotes(27) as the validation set and retrained the PhageBoost model after omitting these genomes from the training data. We benchmarked this model against three prediction tools VIBRANT(8), VirSorter(7) and PHASTER(6) using three predefined metrics: number of regions found, sensitivity and positive predictive value (PPV)(6, 28) (online methods, Figure 2A). VIBRANT and PHASTER were the more conservative when making predictions and had the highest PPV values of 0.84 and 0.69 respectively, but they also found the least amount of the validated phages. PhageBoost with a PPV of 0.44 was higher than VirSorter with 0.31. Given that the validation data genomes could have more than the reported prophages (6), identifying these prophages would negatively influence the PPV. PhageBoost identified the most validated prophage regions with 231, whereas PHASTER identified 224, VirSorter 221 and VIBRANT 192. While PhageBoost reached a higher sensitivity (0.82) than PHASTER (0.78) and VIBRANT (0.73), VirSorter (0.85) had the highest sensitivity finding larger proportions of the validated regions (Figure 2A).

#### Superimposition of TARA Ocean viral samples on single-cell genomic marine microbes

As there are no universal conserved marker genes for phages, we utilized recently published marine datasets(29, 30) to demonstrate that PhageBoost can discover previously unseen prophage signals. We reason that prophage regions in marine bacteria will have similarities to marine phages and by superimposing marine phage sequences on marine bacterial genomes, we will be able to enrich for prophage regions without sequence similarities to existing databases.

We mapped the sequence reads from 223 metagenomic marine viral fractions samples (viromes) from 65 different TARA oceans stations(29) to 5,537 single amplified marine microbial genomes (SAGs)(30). This gave a total of 22,659 unique regions with a signal out of 236,405 contigs (Tables S4-S5) that were consistently detected by at least 1x coverage. We then filtered regions that were at least 10kb long and located 10kb away from the edges, resulting in 746 regions which we hypothesize could be prophage regions within the marine microbial genomes.

Thereafter, we predicted viral signals from all 5,537 SAGs using PhageBoost and three other leading prediction tools - PHASTER, VirSorter and VIBRANT (Table 1). The run times varied between the different programs where PhageBoost was significantly the fastest. PhageBoost finished the prediction in ∼33.5 hours and was 9x faster than VIBRANT, and ∼19x faster than VirSorter with PHASTER taking more than two weeks to get results back from the server (Table 1). The number of predictions also varied widely, with PhageBoost predicting the highest numbers of virome mapped regions overall and for the potential prophage regions, with VirSorter second, and VIBRANT and PHASTER predicting significantly fewer hits mostly finding some of the regions that were found in multiple virome samples (Figure 2B), which could indicate that these regions have already been deposited to virome databases. VirSorter was often found assigning the whole contig as phage, whereas PhageBoost classified smaller regions.

**Table 1.** Prediction software comparisons for the marine SAGs.

|  | Viral regions | Contigs | SAGs<sup>c</sup> | Time per SAG (mins) | Total time (hours) |
| --- | --- | --- | --- | --- | --- |
| PhageBoost | 5757 | 5081 | 2539 | 0.36 ± 0.2 | 33.37 |
| PHASTER | 2402 | 2303 | 1905 | -<sup>a</sup> | 401 |
| VirSorter | 3889 | 3872 | 2552 | 6.8 ± 4.9 | 627.21 |
| VIBRANT | 943 | 942 | 715 | 3.3 ± 1.5 | 301.8<sup>b</sup> |
| Total |  | 234593 | 5537 |  |  |

<sup>a</sup> jobs to webserver submissions took 2 hours 2020, Feb 10 20:02 - 22:00, results received 2020, Feb 13 - Feb 27 13:20; <sup>b</sup> the elapsed time for 5515 genomes, the remaining 242 failed to finish; <sup>c</sup> Single-amplified genomes.

By investigating the sampling locations of the SAGs where a potential prophage region was found, we further observed a substantial increase in predictions making it ecologically significant. The predicted prophage fragments are spread around to multiple locations in the ocean thereby increasing the ecological phage space. (Figure 3 and online methods). As the homology-based approaches cannot go beyond the current known sequence space, our results show that by utilizing the biological feature space and machine learning, PhageBoost is able to generalize and detect previously unknown viral signals for novel hosts such as Prochlorococcus and SAR11. Furthermore, as the viromes’ ecological signal doesn’t get saturated with positive predictions, this suggests that a repository of new prophages awaits to be discovered.

In conclusion, PhageBoost is a fast prophage predictor, which is independent of sequence similarity, able to generalize and therefore is able to predict from new unseen data to facilitate the discovery of previously unknown prophages. We have applied PhageBoost on 5,537 single-cell genomics data and have found significantly more viral regions and considerably faster than with the current state-of-art tools. This finding was validated with available marine virome data. In order to support larger sequencing projects, PhageBoost is freely available as a command-line tool and an interactive online prediction server. This allows future work for inferring the state of lysogenic activity as well as provides new approaches to study the phylogeny of phages and host-phage interactions.

## DATA AVAILABILITY

The PhageBoost predictor is available as an online prediction server at http://www.phageboost.ml and freely available to academic users at GitHub: https://github.com/ku-cbd/PhageBoost. The PB13994 training datasets are available at a frozen archive as https://doi.org/10.17894/ucph.64136536-6353-430b-96ca-701ce89921c4.

The PhageBoost source code is available at GitHub: https://github.com/ku-cbd/PhageBoost.

## SUPPLEMENTARY DATA

Further supplementary data has been deposited at https://doi.org/10.17894/ucph.64136536-6353-430b-96ca-701ce89921c4.

## ACKNOWLEDGEMENT

We thank Tom Delmont for his valuable advice leading to the choice of the marine validation dataset, and Antonio Fernandez Guerra for helpful discussions in taming the possible model bias arising from taxonomy.

