## Supplementary Table and Figure Legends for "Rapid discovery of novel prophages using biological feature engineering and machine learning"

**SUPPLEMENTARY TABLE AND FIGURES LEGENDS**

Table S1. Biological features.

Table S2. Training genomes.

Table S3. Averaged feature importances from training data.

Table S4. Virome regions.

Table S5 Virome alignment rates.

Figure S1. Feature contribution to model output. See Table S3 for the mean absolute Shapley values and Table S1 for feature name explanations.
