## Supplementary figures and images for "Rapid discovery of novel prophages using biological feature engineering and machine learning"

### Figure S1

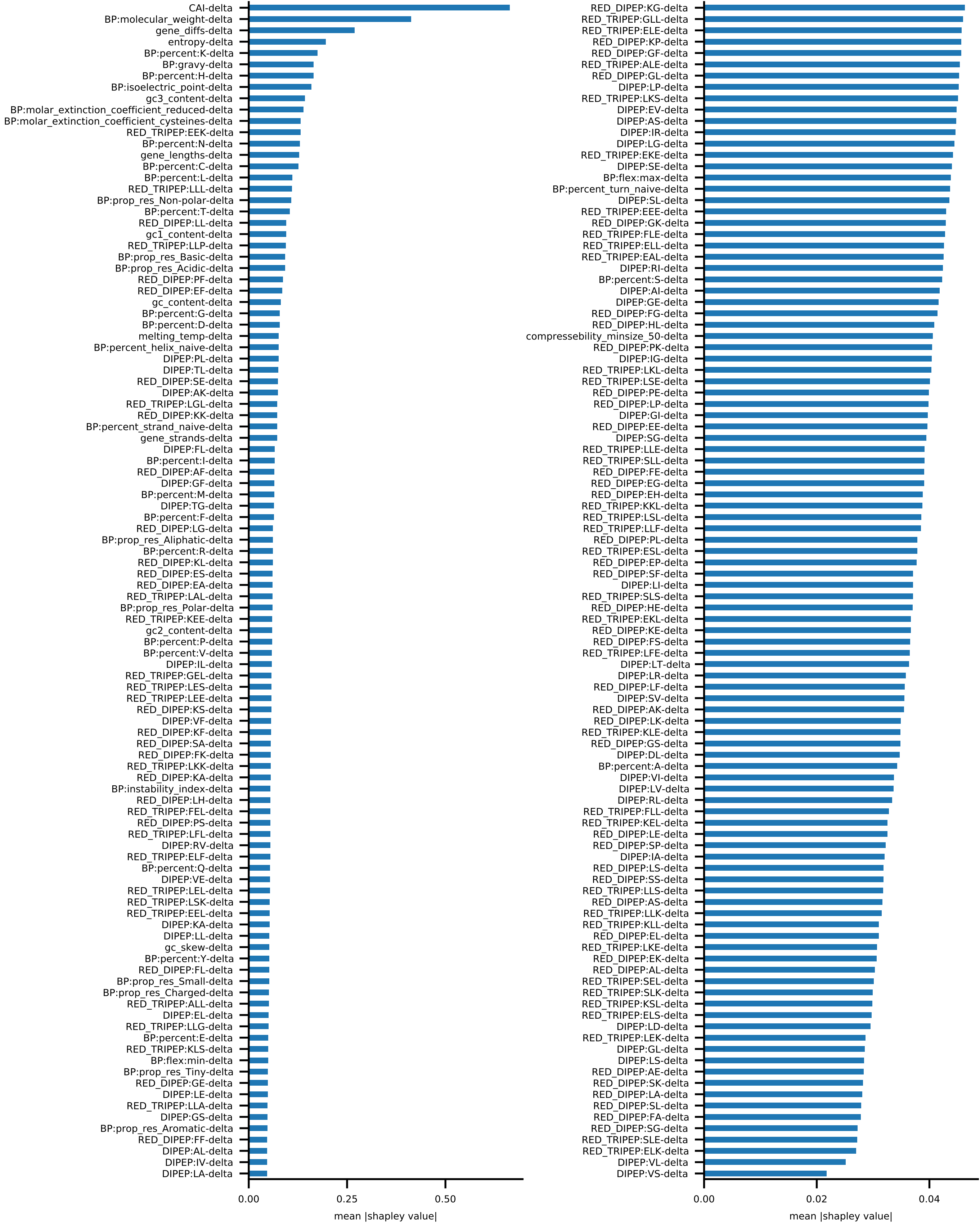
